# The *Mycobacterium tuberculosis* Phosphate-Sensing Pst/SenX3-RegX3 System Regulates ESX-5 Secretion to Evade Host Immunity

**DOI:** 10.1101/400960

**Authors:** Sarah R. Elliott, Dylan W. White, Anna D. Tischler

## Abstract

The *Mycobacterium tuberculosis* Type VII secretion system ESX-5, which has been implicated in virulence, is activated at the transcriptional level by the phosphate starvation responsive Pst/SenX3-RegX3 signal transduction system. Deletion of *pstA1*, which encodes a Pst phosphate transporter component, causes constitutive activation of the response regulator RegX3, hyper-secretion of ESX-5 substrates and attenuation in the mouse infection model. We hypothesized that constitutive activation of ESX-5 secretion causes attenuation of the Δ*pstA1* mutant. To test this, we uncoupled ESX-5 from regulation by RegX3. Using electrophoretic mobility shift assays, we defined a RegX3 binding site in the *esx-5* locus. Deletion or mutation of the RegX3 binding site reversed hyper-secretion of the ESX-5 substrate EsxN by the Δ*pstA1* mutant and abrogated induction of EsxN secretion in response to phosphate limitation by wild-type *M. tuberculosis*. Deletion of the *esx-5* RegX3 binding site (Δ*BS*) suppressed attenuation of the Δ*pstA1* mutant in Irgm1^-/-^ mice, suggesting that constitutive ESX-5 secretion limits *M. tuberculosis* evasion of host immune responses that are independent of Irgm1. However, the Δ*pstA1*Δ*BS* mutant remained attenuated in both NOS2^-/-^ and C57BL/6 mice, suggesting that factors other than ESX-5 secretion also contribute to attenuation of the Δ*pstA1* mutant. In addition, a Δ*pstA1*Δ*esxN* mutant lacking the hyper-secreted ESX-5 substrate EsxN remained attenuated in Irgm1^-/-^ mice, suggesting that ESX-5 substrates other than EsxN cause increased susceptibility to host immunity. Our data indicate that while *M. tuberculosis* requires ESX-5 for virulence, it tightly controls secretion of ESX-5 substrates to avoid elimination by host immune responses.

## INTRODUCTION

Pathogenic bacteria often regulate the activity of specialized protein secretion systems that are required for virulence to ensure release of secreted effectors only at the appropriate stage of infection. Tight control of secretion system activity may limit recognition by the host immune system or prevent expression of complex secretion machines that restrict growth (1, 2). *Mycobacterium tuberculosis*, the causative agent of tuberculosis, encodes five Type VII or ESX specialized protein secretion systems, of which ESX-1, ESX-3 and ESX-5 have been shown to promote pathogenesis (3). *M. tuberculosis* regulates activity of each of these secretion systems in response to signals encountered in the host. Iron limitation activates ESX-3 (4), which plays a role in both iron scavenging and inhibiting phagosome maturation (5, 6). ESX-1 permeabilizes the phagosomal membrane to allow bacterial access to the host cell cytoplasm (7-9). ESX-1 secretion is regulated by two signal transduction systems, PhoPR and MprAB, that respond to acidic pH and cell wall stress, respectively, signals that *M. tuberculosis* encounters in the phagosome (10-13). We recently demonstrated that *M. tuberculosis* activates ESX-5 secretion in response to inorganic phosphate (P_i_) limitation (14). RegX3, a response regulator activated during P_i_ limitation, directly activates transcription of a subset of *esx-5* genes leading to increased production of ESX-5 secretion system core components and enhanced secretion of the EsxN and PPE41 substrates (14).

Though the precise function of ESX-5 remains unclear, it appears to influence nutrient acquisition to enable *M. tuberculosis* replication (15-17) and to promote host cell necrosis by activating the inflammasome and stimulating IL-1β secretion (18, 19). In the related pathogen *Mycobacterium marinum*, ESX-5 secretes most proteins that belong to the mycobacteria-specific PE and PPE protein families (16, 20). The *M. tuberculosis* PE and PPE proteins are strongly immunogenic in mice; immune responses to PE and PPE antigens depend on a functional ESX-5 secretion system, suggesting that *M. tuberculosis* also secretes many PE and PPE proteins via ESX-5 (21). ESX-5 is also likely to be active during infection since T cells specific for the ESX-5 substrate EsxN have been detected in humans with latent tuberculosis (22, 23).

Activation of the RegX3 response regulator and induction of ESX-5 secretion is inhibited during growth in P_i_-replete conditions by the Pst P_i_ uptake system (24). Deletion of *pstA1*, which encodes a Pst system trans-membrane component, causes constitutive activation of RegX3, constitutive expression of *esx-5* genes, and hyper-secretion of ESX-5 substrates, independent of P_i_ availability (14). We previously demonstrated that a Δ*pstA1* mutant is attenuated during the chronic phase of infection in wild-type C57BL/6 mice and exhibits strongly reduced replication and virulence in two immune-deficient strains of mice, NOS2^-/-^ and Irgm1^-/-^, that fail to control infection with wild-type *M. tuberculosis* (24). NOS2^-/-^ mice lack the interferon-gamma (IFN-γ) inducible nitric oxide synthase that generates toxic reactive nitrogen species (25). Although NOS2^-/-^ mice are assumed to have a cell-intrinsic defect in their ability to control *M. tuberculosis* replication (26), they also fail to inhibit neutrophil recruitment to the lung, which creates a nutrient-rich environment that enhances *M. tuberculosis* replication (27, 28). Irgm1 encodes an IFN-γ inducible GTPase that was originally described to restrict *M. tuberculosis* replication in a cell-intrinsic manner by mediating phagosome acidification, possibly via induction of autophagy (29, 30). However, Irgm1 is also required for hematopoietic stem cell renewal (31); Irgm1^-/-^ mice become leukopenic upon infection with intracellular pathogens, including mycobacteria (32), which also likely contributes to their profound susceptibility to infection. We previously demonstrated that attenuation of the Δ*pstA1* mutant in NOS2^-/-^ mice was due to the constitutive activation of RegX3; a Δ*pstA1*Δ*regX3* double mutant progressively replicated in the lungs and caused death of the animals (24). It remains unclear whether constitutive activation of RegX3 similarly contributes to attenuation of the Δ*pstA1* mutant in either Irgm1^-/-^ or C57BL/6 mice, because a Δ*regX3* single mutant was also attenuated in these mouse strains (24).

We hypothesized that constitutive activation of *esx-5* transcription and hyper-secretion of ESX-5 substrates driven by constitutively activated RegX3 causes virulence attenuation of the Δ*pstA1* mutant. *M. tuberculosis* requires ESX-5 for replication *in vitro* (15, 33), so we were unable to construct mutants lacking ESX-5 function to test this possibility. Instead, we took a targeted approach to uncouple ESX-5 from regulation by RegX3. We defined the RegX3 binding site in the *esx-5* locus and generated targeted mutations that disrupt RegX3 binding. Mutation of the RegX3 binding site prevented induction of *esx-5* gene expression and ESX-5 secretion during P_i_ limitation by wild-type *M. tuberculosis*, and reversed the over-expression of *esx-5* genes and hyper-secretion the ESX-5 substrate EsxN by the Δ*pstA1* mutant. Deletion of the *esx-5* RegX3 binding site also suppressed attenuation of the Δ*pstA1* mutant specifically in Irgm1^-/-^ mice. Our results suggest hyper-secretion of ESX-5 substrates sensitizes *M. tuberculosis* to a host immune response that is independent of Irgm1 and that *M. tuberculosis* regulates ESX-5 secretion in response to P_i_ availability in the host to evade this host immune response.

## RESULTS

### Defining a RegX3 binding site in the *esx-5* locus

We previously demonstrated that RegX3 directly regulates ESX-5 activity at the transcriptional level via binding to a 125 bp sequence within the *ppe27-pe19* intergenic region in the *esx-5* locus (Fig. 1A) (14). RegX3 was not included in a prior study that mapped the binding sites of most *M. tuberculosis* transcription factors (34), so a RegX3 consensus binding sequence has yet to be described. To more precisely define the *esx-5* RegX3 binding site, we conducted competitive electrophoretic mobility shift assays (EMSAs) using purified recombinant His_6_-RegX3. Our previous work demonstrated that RegX3 binds within the sequence −151 to −27 bp relative to the *pe19* start codon (Probe A, Fig. 1A and 1B) (14). Binding reactions including excess unlabeled competitors comprising the 5’ (−151 to −91) or 3’ (−90 to −28) halves of Probe A demonstrated that RegX3 binds to the 5’ region; only addition of the 5’ competitor resulted in reversal of the mobility shift (Fig. 1B). These data indicate that RegX3 binds within −151 to −91 bp relative to the *pe19* start codon.

**Figure 1.**
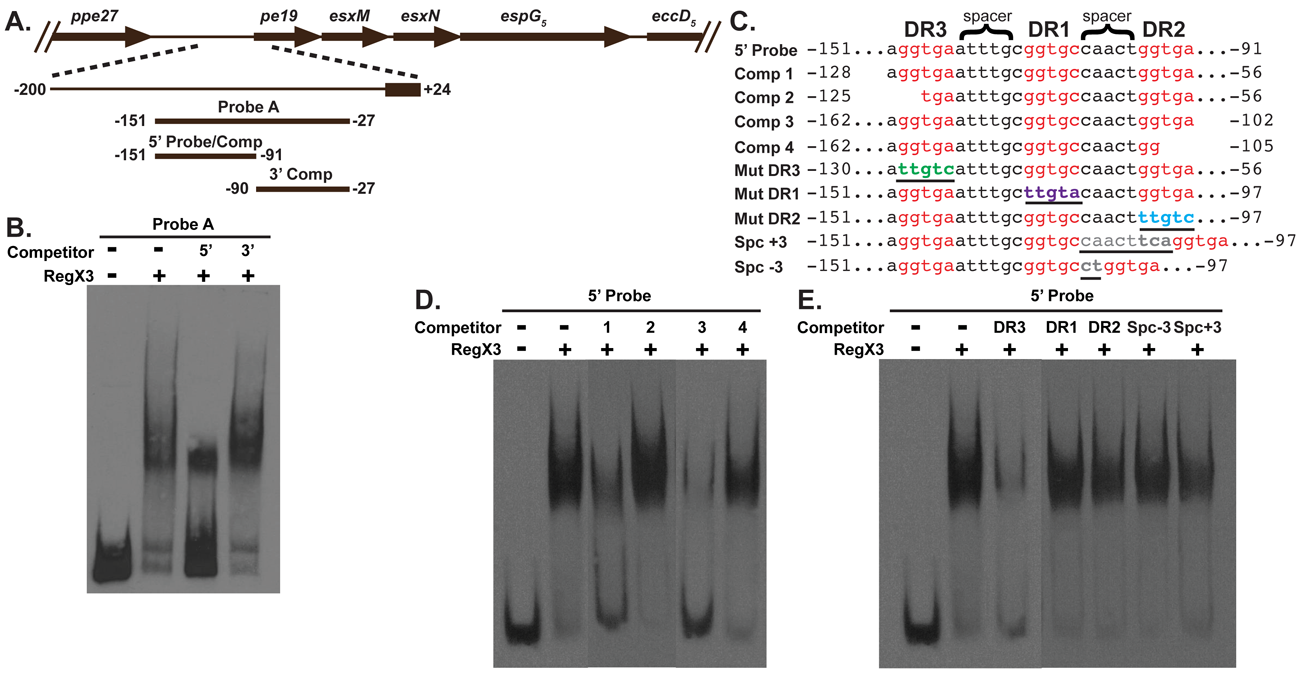
Competitive EMSAs define a RegX3 binding site 5’ of *pe19* in the *esx-5* locus. (A) Schematic depicting the location of EMSA probes and competitors in the *esx-5* locus. Positions of the 5’ and 3’ ends of probe and competitor sequences relative to the *pe19* translational start site are indicated. (C) Sequences of the 5’ Probe, truncated competitors that defined the 5’ and 3’ ends of the RegX3 binding site, and mutated competitors that defined sequence elements required for RegX3 binding. Direct repeats (red text) and the 5’ and 3’ ends of each competitor relative to the *pe19* translational start site are indicated. Mutated sequences are highlighted by underlines and green (DR3), purple (DR1), blue (DR2) or gray (spacer) text. (B, D, and E) EMSA analysis of binding between purified His_6_-RegX3 (0.5 μg), DIG-labeled probe (0.5 ng), and unlabeled competitors (200 ng), as indicated. Results are representative of two independent experiments.

To further define the *esx-5* RegX3 binding site, we performed additional competitive EMSAs using the 61 bp −151 to −91 segment as the labeled probe (Fig. 1A and 1C, 5’ Probe) and a series of unlabeled competitors that truncate the 5’ Probe sequence at either the 5’ or 3’ end, added in excess. A complete list of competitors tested and their ability to compete with the 5’ Probe for RegX3 binding is provided in Table S1. Competitors that defined the 5’ and 3’ ends of the RegX3 binding site are shown (Fig. 1C). Excess unlabeled Competitor 1, which truncates the 5’ Probe at the 5’ end, reversed the mobility shift, indicating that RegX3 binds to this competitor (Fig. 1C and 1D). However, RegX3 did not bind Competitor 2, which truncates an additional three bp at the 5’ end, since the mobility shift was unperturbed (Fig. 1C and 1D), indicating that one or more base pairs removed from Competitor 2 are essential for RegX3 binding. Therefore, the 5’ end of the RegX3 binding site is located near position −128 relative to the *pe19* start codon. Similarly for the 3’ end, excess Competitor 3 reversed the mobility shift, indicating RegX3 can bind to this sequence, but excess Competitor 4, which eliminates an additional three bp from the 3’ end, failed to alter the mobility shift (Fig. 1C and 1D). These data demonstrate that the three bp removed from Competitor 4 relative to Competitor 3 are required for RegX3 binding, and thus define the 3’ end of the RegX3 binding site at −102 relative to the *pe19* start codon. Collectively, our data indicate that RegX3 binds to a 27 bp sequence located at −128 to −102 relative to the *pe19* start codon in the *esx-5* locus.

### Defining essential sequence elements for RegX3 binding *in vitro*

RegX3 is a member of the OmpR/PhoB family of winged helix-turn-helix response regulators that typically bind to direct repeat DNA sequences (35). We previously identified an imperfect direct repeat separated by a 5 bp spacer in the 5’ Probe sequence (DR1 and DR2, Fig. 1C) (14). Further examination revealed a third imperfect direct repeat (DR3) 5’ of the first two and separated from DR1 by a 6 bp spacer (Fig. 1C). All three direct repeats are contained within the −128 to −102 region relative to the *pe19* start codon. To determine if these sequence elements are required for RegX3 binding, EMSAs were performed using competitor DNA harboring mutations in the individual direct repeats or spacer elements. For each direct repeat element, all five bp of the direct repeat were altered by transversion (Fig. 1C). We altered the spacer sequence between DR1 and DR2 by either adding or removing three base pairs (Spc+3 and Spc-3, respectively, Fig. 1C). Each mutated unlabeled competitor was tested for the ability to compete with the 5’ probe for binding to RegX3 when added in excess. The mutated DR3 competitor reversed the mobility shift, indicating RegX3 can still bind this sequence (Fig. 1E). However, the mutated DR1 or DR2 competitors both failed to reverse the mobility shift, indicating that RegX3 cannot bind these mutated sequences (Fig. 1E). These data indicate that the DR1 and DR2 sequence elements are required for RegX3 binding *in vitro*. Altering the spacing between DR1 and DR2, either by adding or removing 3 bp, abrogates RegX3 binding, since the Spc+3 and Spc-3 competitors also failed to reverse the mobility shift (Fig. 1E). This indicates that RegX3 requires a 5 bp spacer between DR1 and DR2 for *in vitro* binding. The 27 bp RegX3 binding site sequence, including DR1 and DR2, located approximately 100 bp upstream of the *pe19* start codon is consistent with RegX3 functioning as a transcriptional activator of *esx-5* genes (14).

### RegX3 binding site mutations in the Δ*pstA1* mutant reverse *esx-5* over-expression and hyper-secretion of EsxN

We previously demonstrated that *esx-5* transcripts are over-expressed during growth in P_i_-replete conditions in the Δ*pstA1* mutant due to constitutive activation of RegX3 (14). To determine if *esx-5* over-expression also depends upon the *esx-5* RegX3 binding site that we defined, we introduced three distinct RegX3 binding site mutations at the intergenic region 5’ of *pe19* on the chromosome of the *M. tuberculosis* Δ*pstA1* mutant. The DR2 direct repeat mutant (Δ*pstA1*_DR2_) harbors the transversion mutations in DR2 identical those tested for RegX3 binding *in vitro* (Fig. 1C). The spacer mutant (Δ*pstA1*_Spc+3_) harbors three additional bp between DR1 and DR2, identical to the Spc+3 mutation tested for RegX3 binding *in vitro* (Fig. 1C). Finally, the binding site deletion mutant (Δ*pstA1*Δ*BS*) harbors a deletion of the complete 27 bp RegX3 binding site located at −128 to −102 bp relative to the *pe19* start codon. We tested expression of *esx-5* genes in the Δ*pstA1*_DR2_, Δ*pstA1*_Spc+3_ and Δ*pstA1*Δ*BS* binding site mutants grown in standard P_i_-rich medium (Fig. 2A). The Δ*pstA1* mutant exhibited significant over-expression of the *pe19* and *espG*_*5*_ transcripts (*P*<0.0001) and more than 3-fold over-expression of *eccD*_*5*_ as compared to the WT control (Fig. 2A). As previously reported, over-expression of these transcripts was dependent on RegX3 since expression of each gene was restored to the WT level in the Δ*pstA1*Δ*regX3* mutant (Fig. 2A) (14). In both the Δ*pstA1*_DR2_ and Δ*pstA1*Δ*BS* mutants, transcription of *pe19, espG*_*5*_, and *eccD*_*5*_ was similarly restored to levels that were nearly the same as and not significantly different from the WT control (Fig. 2A). Both the Δ*pstA1*_DR2_ and Δ*pstA1*Δ*BS* mutants also exhibited statistically significant reductions in *pe19* and *espG*_*5*_ transcription relative to the Δ*pstA1* parental control (Fig. 2A). The *pe19, espG*_*5*_, and *eccD*_*5*_ transcripts were detected at intermediate levels in the Δ*pstA1*_Spc+3_ mutant that were not significantly reduced as compared to the Δ*pstA1* parental strain (Fig. 2A). These data demonstrate that the RegX3 binding site within the *esx-5* locus, and the DR2 sequence in particular, is required for RegX3-mediated over-expression of *esx-5* genes in the Δ*pstA1* mutant.

**Figure 2.**
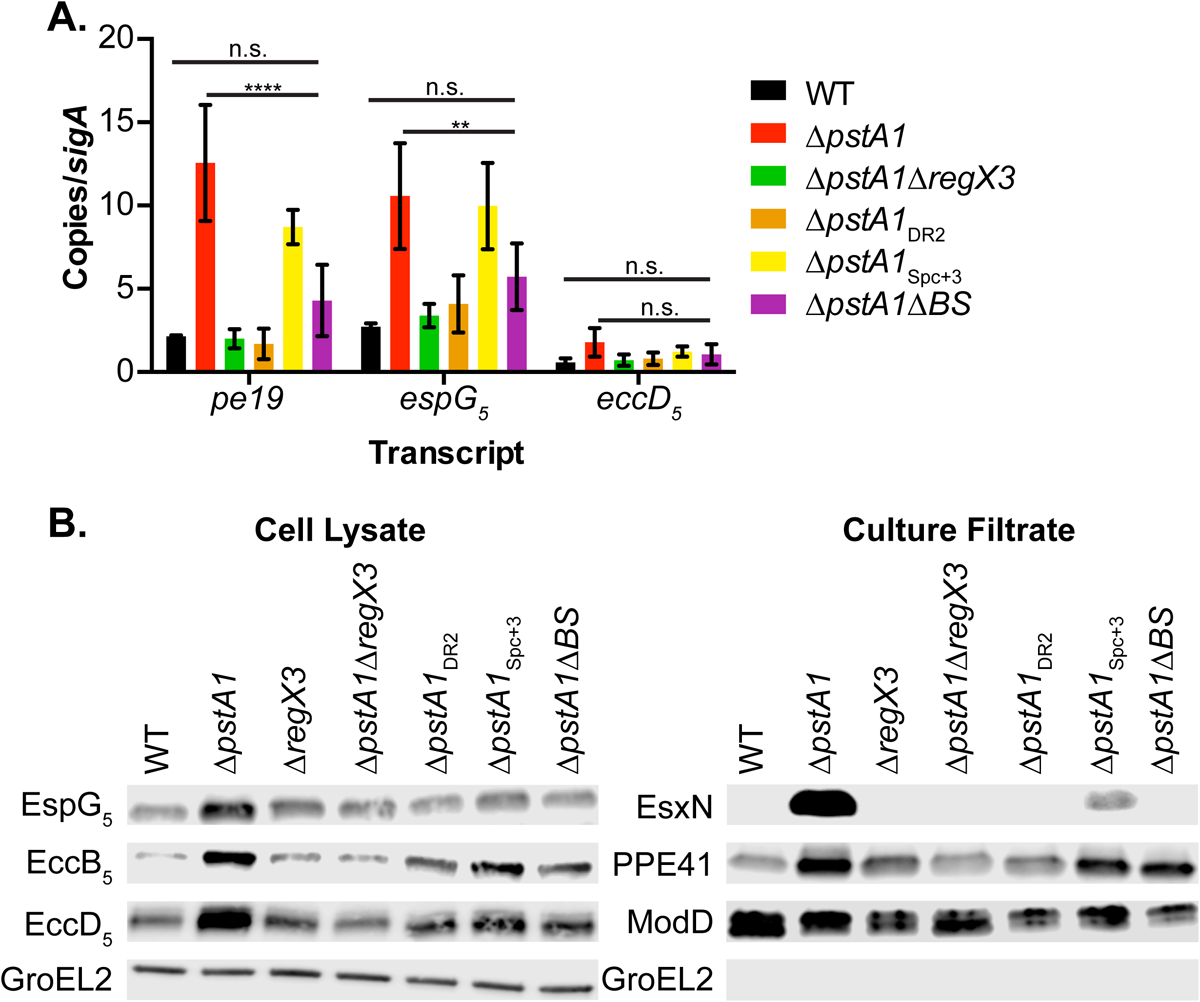
Mutation of the *esx-5* RegX3 binding site suppresses over-expression of *esx-5* genes and hyper-secretion of EsxN by the Δ*pstA1* mutant. (A) Transcript abundance of *pe19, espG*_*5*_ and *eccD*_*5*_ relative to *sigA* were determined by quantitative RT-PCR for the indicated strains grown to mid-logarithmic phase in 7H9 complete medium. Results are the means ± standard deviations of three independent experiments. \*\**P*<0.01, \*\*\*\**P*<0.0001. (B) The indicated strains were grown in Sauton’s medium without Tween-80. Cell lysates (10 μg) and culture filtrates (5 μg) were separated and analyzed by Western blotting to detect the indicated proteins. Results shown are from a single experiment and are representative of two independent experiments.

RegX3 is a global response regulator that activates and represses many genes outside of the *esx-5* locus (24). To determine if the RegX3 binding site mutations that we introduced perturbed regulation exclusively at the *esx-5* locus, we examined transcription of other genes that are over-expressed by the Δ*pstA1* mutant in a RegX3-dependent manner, but that are not associated with *esx-5* (24). The *udgA* and *mgtA* transcripts were over-expressed by the Δ*pstA1* mutant relative to both the WT and Δ*pstA1*Δ*regX3* strains (Fig. S1). Both *udgA* and *mgtA* transcripts remained significantly over-expressed in the Δ*pstA1*_DR2_, *ΔpstA1*_Spc+3_ and Δ*pstA1*Δ*BS* mutants (Fig. S1). These data demonstrate that mutation of the RegX3 binding site sequence within the *esx-5* locus does not globally alter RegX3 activity.

To determine if the decreased transcription of *esx-5* genes in the RegX3 binding site mutants translates to changes in stability or activity of the ESX-5 secretion system, we monitored production of ESX-5 conserved components and secretion of the ESX-5 substrates EsxN and PPE41 by the Δ*pstA1* RegX3 binding site mutants. We observed hyper-secretion of the ESX-5 substrates EsxN and PPE41 and over-production of the cytosolic ESX-5 chaperone EspG_5_ and ESX-5 secretion machinery components EccB_5_ and EccD_5_ by the Δ*pstA1* mutant relative to the WT control (Fig. 2B). This response required RegX3 (Fig. 2B), consistent with our prior report (14). We detected reduced amounts of the EspG_5_, EccB_5_ and EccD_5_ proteins in all three Δ*pstA1* RegX3 binding site mutants as compared to the Δ*pstA1* mutant (Fig. 2B). EsxN hyper-secretion was reversed in both the Δ*pstA1*_DR2_ and Δ*pstA1*Δ*BS* mutants, reaching levels that were undetectable, comparable to both the WT and Δ*pstA1*Δ*regX3* mutant controls (Fig. 2B). We detected EsxN secretion by the Δ*pstA1*_Spc+3_ mutant but at a 6-fold reduced abundance as compared to the Δ*pstA1* mutant (Fig. 2B). Secretion of PPE41 was also decreased in each of the RegX3 binding site mutants relative to the Δ*pstA1* parental strain, but remained approximately 2-fold increased as compared to the WT control (Fig. 2B). It is possible either that RegX3 controls PPE41 secretion by a mechanism independent of its regulation of *esx-5* transcription, or that decreased secretion of EsxN frees the ESX-5 secretion apparatus to translocate other substrates including PPE41. The ModD control confirmed equivalent loading of the culture filtrate fraction; the GroEL2 control confirmed equivalent loading of the cell lysate fraction and demonstrated that cell lysis did not contaminate the culture filtrate (Fig. 2B). These results indicate that the RegX3 binding site in the *esx-5* locus is required for the over-production of ESX-5 secretion system core components and hyper-secretion of EsxN by the Δ*pstA1* mutant.

### Mutation of the RegX3 binding site in the *esx-5* locus prevents ESX-5 induction during phosphate limitation

We previously demonstrated that P_i_ limitation triggers ESX-5 activity in WT *M. tuberculosis*, and that this response requires RegX3 (14). To determine if the RegX3 binding site is also required for induction of *esx-5* transcription in response to P_i_ limitation, we generated a strain lacking the RegX3 binding site in the WT Erdman strain background (Δ*BS*) and conducted qRT-PCR experiments to monitor *esx-5* gene expression. The WT, Δ*regX3* and Δ*BS* strains were grown in either P_i_-free medium or P_i_-replete medium as a control. In P_i_-replete conditions, *esx-5* transcripts were expressed at a basal level in all of the strains (Fig. 3B). Statistically significant increases in *pe19* and *espG*_*5*_ transcription were detected for the Δ*BS* mutant, but the changes were less than 1.5-fold (Fig. 3B). The *pe19, espG*_*5*_ and *eccD*_*5*_ transcripts were induced 7.9, 4.9, and 3.7-fold, respectively, by WT *M. tuberculosis* during growth in P_i_-free medium relative to the P_i_-replete control (Fig. 3A). The Δ*regX3* mutant failed to induce *pe19, espG*_*5*_ or *eccD*_*5*_ transcription in response to P_i_ limitation, consistent with our previous reports (Fig. 3A) (14, 36). The Δ*BS* mutant similarly failed to induce *pe19, espG*_*5*_ or *eccD*_*5*_ transcription in response to P_i_ limitation (Fig. 3A); the level of each transcript was significantly different from that of the WT control and not significantly different from that of the Δ*regX3* mutant (Fig. 3A). These data demonstrate that the RegX3 binding site in the *esx-5* locus is required for activation of *esx-5* transcription in response to P_i_ limitation.

**Figure 3.**
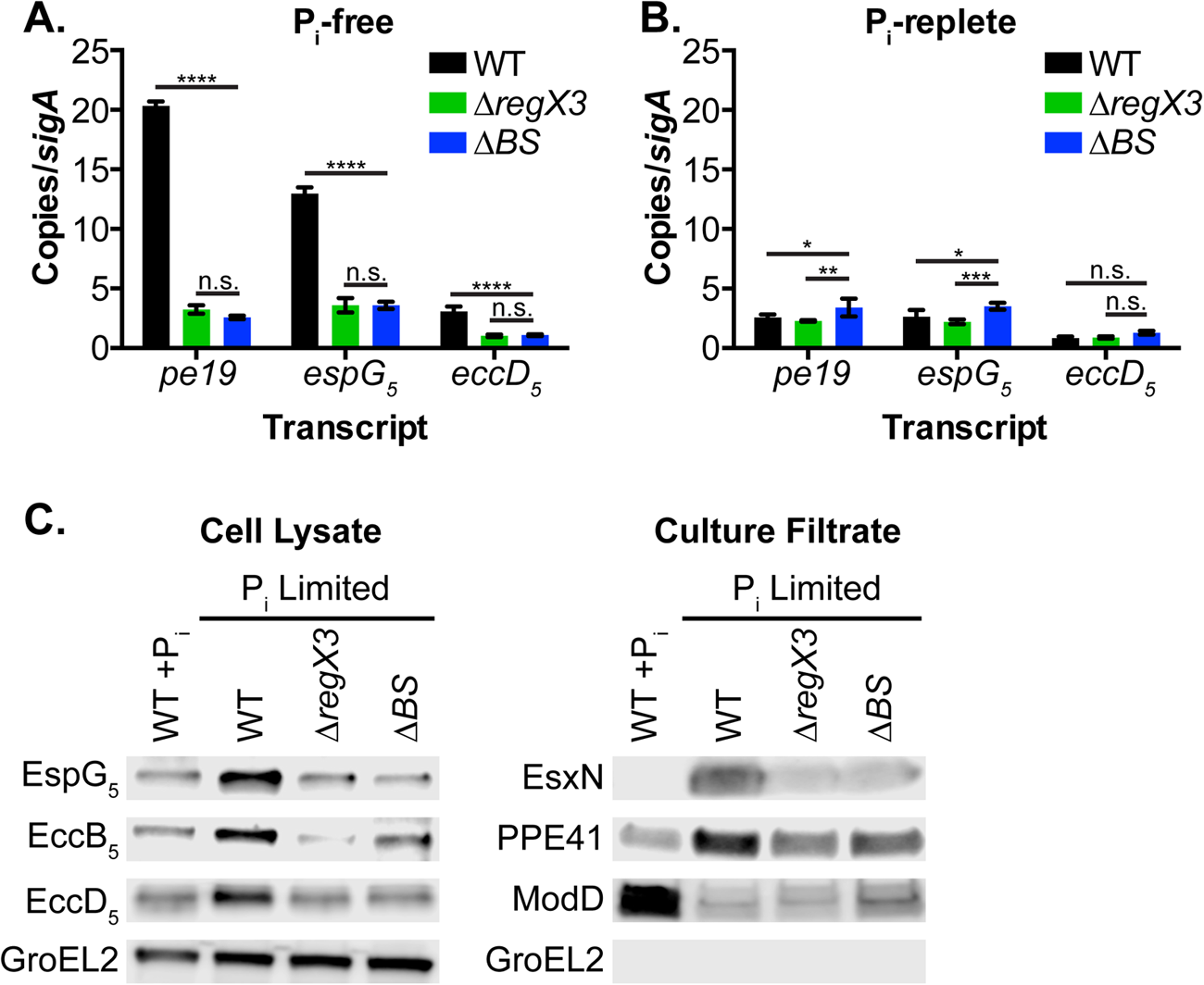
Deletion of the *esx-5* RegX3 binding site prevents activation of ESX-5 secretion in response to P_i_ limitation. (A & B) Transcript abundance of *pe19, espG*_*5*_ and *eccD*_*5*_ relative to *sigA* were determined by quantitative RT-PCR for the indicated strains grown to mid-logarithmic phase in P_i_-free 7H9 medium (A) or P_i_-replete 7H9 complete medium (B). Results are the means ± standard deviations of three independent experiments. \**P*<0.05, \*\**P*<0.01,\*\*\**P*<0.0001, \*\*\*\**P*<0.0001. (C) The indicated strains were grown in Sauton’s medium without Tween-80 (WT +P_i_) or in P_i_-limiting (2.5 μM P_i_) Sauton’s medium without Tween-80. Cell lysates (10 μg) and culture filtrates (5 μg) were separated and analyzed by Western blotting to detect the indicated proteins. Results shown are from a single experiment and are representative of two independent experiments.

We evaluated production of ESX-5 conserved components and secretion of the ESX-5 substrates EsxN and PPE41 during P_i_ limitation in the Δ*BS* mutant by Western blotting. Production of EspG_5_, EccB_5_ and EccD_5_ was induced in WT *M. tuberculosis* during P_i_ limitation (Fig. 3C), as previously demonstrated (14). Increased production or stability of EspG_5_, EccB_5_ and EccD_5_ during P_i_ limitation was abrogated in both the Δ*regX3* and Δ*BS* mutants (Fig. 3C). The GroEL2 control confirmed equivalent loading of cell lysate proteins (Fig. 3C). Secretion of EsxN and PPE41 was induced in the WT strain during P_i_ limitation, as previously reported (14) (Fig. 3C). Induction of EsxN secretion during P_i_ limitation was prevented by either the Δ*regX3* or the Δ*BS* mutation (Fig. 3C). The Δ*regX3* mutant also exhibited modestly reduced PPE41 secretion during P_i_ limitation, as previously reported (14) (Fig. 3C). In contrast, the Δ*BS* mutant induced PPE41 secretion during P_i_ limitation similarly to the WT control (Fig. 3C), consistent with our results demonstrating intermediate PPE41 secretion by the Δ*pstA1*Δ*BS* mutant. The ModD control confirmed equivalent loading of the P_i_-limited culture filtrate fraction (Fig. 3C); the decreased secretion of ModD during P_i_ limitation relative to the P_i_-replete control (Fig. 3C) was consistent with our previous report (14). The GroEL2 control confirmed that cell lysis did not contaminate the culture filtrate (Fig. 3C). Collectively, these data suggest that the RegX3 binding site within the *esx-5* locus is necessary for induction of EsxN secretion during P_i_ limitation.

### The *esx-5* RegX3 binding site deletion suppresses attenuation of the Δ*pstA1* mutant in Irgm1^-/-^ mice

To determine if constitutive hyper-secretion of ESX-5 substrates contributes to attenuation of the Δ*pstA1* mutant we infected C57BL/6, NOS2^-/-^ and Irgm1^-/-^ mice via the aerosol route with ∼100 CFU of WT, Δ*pstA1,* or Δ*pstA1*Δ*BS M. tuberculosis* strains. All Irgm1^-/-^ mice succumbed to infection with WT Erdman by 4 weeks post-infection, and bacterial loads reached over 10^9^ in the lungs (Fig. 4A). Irgm1^-/-^ mice controlled replication of the Δ*pstA1* mutant after 2 weeks post infection (Fig. 4A), consistent with previous results (24). In contrast, the Δ*pstA1*Δ*BS* mutant replicated progressively in the lungs of Irgm1^-/-^ mice. At 4 weeks post-infection, mean bacterial CFU in the lungs of Irgm1^-/-^ mice infected with the Δ*pstA1*Δ*BS* mutant were increased 40-fold as compared to mice infected with the Δ*pstA1* mutant, though this difference did not achieve statistical significance (Fig. 4A). However, by 6 weeks post-infection, the Δ*pstA1*Δ*BS* mutant reached nearly the same final bacterial burden in the lungs of Irgm1^-/-^ mice as the WT control (Fig. 4A). At 6 weeks, the bacterial burden of the Δ*pstA1*Δ*BS* mutant in the lungs was over 1000-fold higher than the Δ*pstA1* mutant, and this difference was statistically significant (Fig. 4A). In these experiments, several of the Irgm1^-/-^ mice infected with the Δ*pstA1*Δ*BS* mutant were moribund at the 6 week time point, while our previous experiments demonstrated that Irgm1^-/-^ mice infected with the Δ*pstA1* mutant all survive for at least 14 weeks (24). These data suggest that attenuation of the Δ*pstA1* mutant in Irgm1^-/-^ mice is due, at least in part, to constitutive activity of the ESX-5 secretion system and increased secretion of one or more ESX-5 substrates. These data further suggest that hyper-secretion of ESX-5 substrates sensitizes *M. tuberculosis* to a host immune response that is independent of Irgm1.

**Figure 4.**
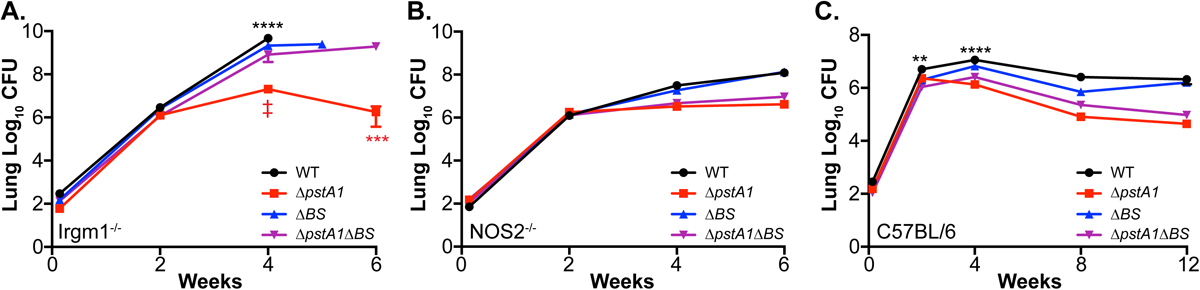
Deletion of the *esx-5* RegX3 binding site in the Δ*pstA1* mutant restores virulence in Irgm1^-/-^ mice. Irgm1^-/-^ (A), NOS2^-/-^ (B) or C57BL/6J (C) mice were infected by the aerosol route with ∼100 CFU of the *M. tuberculosis* WT, Δ*pstA1*, Δ*BS*, or Δ*pstA1*Δ*BS* strain. Groups of mice (*n*=4) were sacrificed at the indicated time points and bacterial CFU were enumerated by plating serial dilutions of lung homogenates. Results are the means ± standard errors of the means. Results for the Δ*pstA1*Δ*BS* mutant in Irgm1^-/-^ mice are from one representative experiment of two independent experiments. All other results are from a single experiment. Data for the WT control in panel B are reproduced from (24) for comparison with the Δ*BS* mutant. Asterisks indicate statistically significant differences between WT and Δ*BS* (black) or between Δ*pstA1* and Δ*pstA1*Δ*BS* (red). \*\**P*<0.01, \*\*\**P*<0.001, \*\*\*\**P*<0.0001,‡*P*=0.1353.

To determine if the modest attenuation of the Δ*pstA1*Δ*BS* mutant relative to the WT control might be due to the Δ*BS* mutation, we performed similar aerosol infection experiments with the Δ*BS* mutant. The Δ*BS* mutant exhibited a modest but statistically significant decrease in lung bacterial burden at the 4 week time point compared to the WT control, but all mice were moribund by the 5 week time point and were euthanized (Fig. 4A). These data suggest that the partially attenuated phenotype of the Δ*pstA1*Δ*BS* mutant in Irgm1^-/-^ mice may be due to an inability to induce ESX-5 secretion in response to P_i_ limitation.

In NOS2^-/-^ mice, in contrast, the Δ*BS* mutation had no statistically significant effect on the ability of either the Δ*pstA1* mutant or WT bacteria to replicate in the lungs (Fig. 4B). These data suggest that other factors besides increased ESX-5 secretion contribute to the attenuation of the Δ*pstA1* mutant in NOS2^-/-^ mice.

Similarly, in C57BL/6 mice, deletion of the *esx-5* RegX3 binding site sequence failed to suppress the attenuated phenotype of the Δ*pstA1* mutant during the chronic phase of infection (Fig. 4C). There were no statistically significant differences in lung bacterial burden between the Δ*pstA1* and Δ*pstA1*Δ*BS* mutants at any time point (Fig. 4C). In addition, at each time point the CFUs in the lungs of C57BL/6 mice infected with either the Δ*pstA1* mutant or the Δ*pstA1*Δ*BS* mutant were significantly different from the WT control (Fig. 4C). To determine if the Δ*BS* mutation causes attenuation, we infected C57BL/6 mice by the aerosol route with the Δ*BS* mutant. We observed modest but statistically significant decreases in lung bacterial burden at the 2 week and 4 week time points in Δ*BS*-infected mice relative to mice infected with the WT control (Fig. 4C). However, by 12 weeks post-infection, CFU in the lungs of both WT- and Δ*BS*-infected mice were similar (Fig. 4C). Taken together, these data suggest that other factors besides increased ESX-5 secretion contribute to attenuation of the Δ*pstA1* mutant during the chronic phase of infection in C57BL/6 mice and that regulation of ESX-5 secretion in response to P_i_ limitation enhances acute phase replication of *M. tuberculosis* in the lungs.

### EsxN hyper-secretion does not cause attenuation of the Δ*pstA1* mutant

To investigate whether attenuation of the Δ*pstA1* mutant is due to inappropriate hyper-secretion of the ESX-5 substrate EsxN specifically, we deleted *esxN* in both WT and Δ*pstA1* mutant backgrounds. The Δ*esxN* deletion was verified by PCR (data not shown) and qRT-PCR; the *esxN* transcript was not detected in either Δ*esxN* mutant (Fig. 5A). Deletion of *esxN* did not alter abundance of the downstream *espG*_*5*_ transcript in either the WT or Δ*pstA1* mutant background (Fig. 5A), suggesting that the Δ*esxN* deletion is not polar. To verify that EsxN is not produced or secreted by the Δ*esxN* and Δ*pstA1*Δ*esxN* mutants, we performed Western blots. While secreted EsxN was not detected in either the WT or Δ*esxN* strains, a protein or proteins that reacted with our anti-EsxN anti-serum was still detected in the secreted fraction of the Δ*pstA1*Δ*esxN* mutant, though at a decreased level compared to the Δ*pstA1* parental control (Fig. 5B). These data suggest that while EsxN itself is hyper-secreted by the Δ*pstA1* mutant, our anti-EsxN anti-serum also detects one or more of the four EsxN paralogs encoded outside the *esx-5* locus that each exhibit >92.5% amino acid sequence identity with EsxN (37). Similar cross-reactivity of anti-EsxN anti-serum was previously described (17). Our data further suggest that secretion of one or more of these EsxN paralogs is also increased in the Δ*pstA1* mutant.

**Figure 5.**
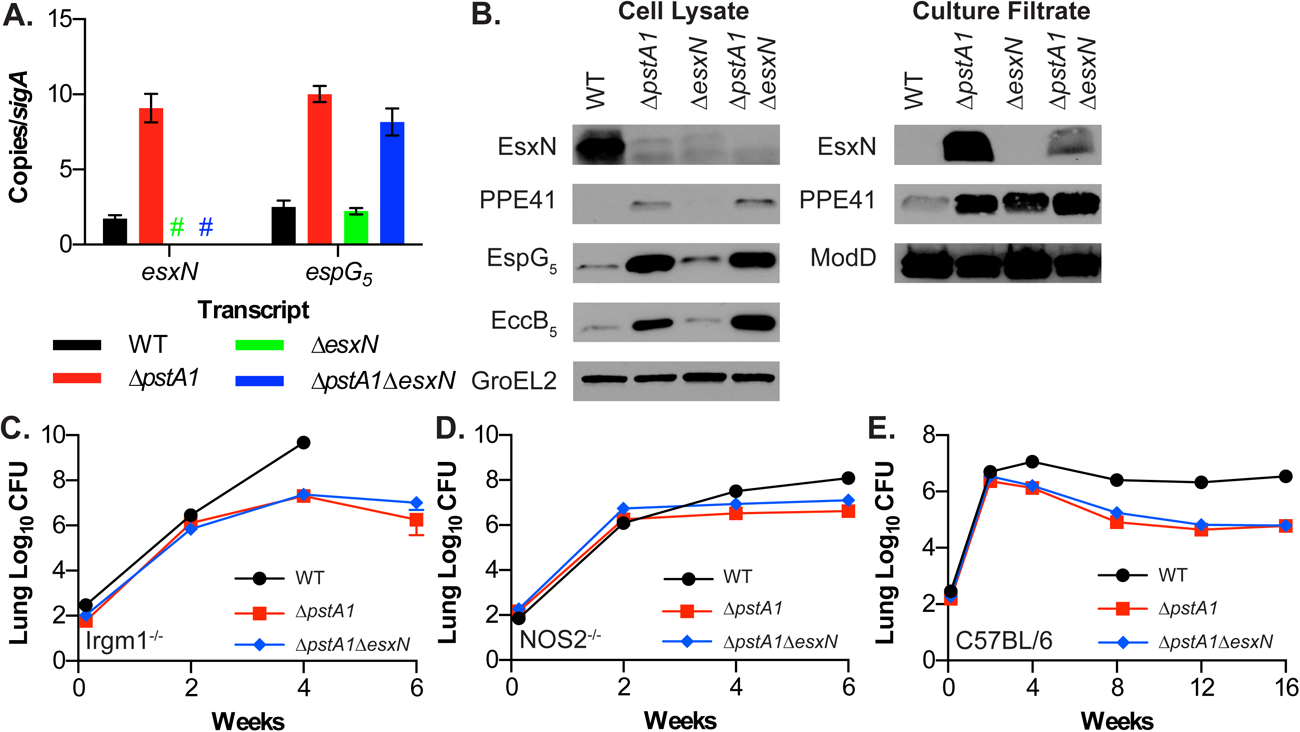
Hyper-secretion of EsxN does not cause attenuation of the Δ*pstA1* mutant. (A) Abundance of the *esxN* and *espG*_*5*_ transcripts relative to *sigA* was determined by quantitative RT-PCR for the indicated strains grown to mid-logarithmic phase in 7H9 complete medium. Results are the means ± standard deviations of three independent experiments. Colored # indicates the transcript was not detected in the corresponding mutant. (B) The indicated strains were grown in Sauton’s medium without Tween-80. Cell lysates (10 μg) and culture filtrates (5 μg) were separated and analyzed by Western blotting to detect the indicated proteins. Results shown are from a single experiment and are representative of two independent experiments. (C-E) Aerosol infection of mice. Irgm1^-/-^ (C), NOS2^-/-^ (D) or C57BL/6J (E) mice were infected with ∼100 CFU of the *M. tuberculosis* WT, Δ*pstA1*, or Δ*pstA1*Δ*esxN* strain. Groups of mice (*n*=4) were sacrificed at the indicated time points and bacterial CFU were enumerated by plating serial dilutions of lung homogenates. Results are the means ± standard errors of the means and are from a single experiment. Data for the Δ*pstA1*Δ*esxN* mutant in Irgm1^-/-^ mice are representative of two independent experiments. Data for the WT and Δ*pstA1* mutant controls are reproduced from Figure 4.

We performed additional Western blots to verify that the Δ*esxN* deletion did not alter production of ESX-5 core components. Because co-dependent secretion of substrates has been observed for the ESX-1 secretion system (38-40), we also examined if deletion of *esxN* altered PPE41 secretion. Deletion of *esxN* in WT *M. tuberculosis* did not change production of the ESX-5 proteins EspG_5_ or EccB_5_, but did cause an increase in PPE41 secretion (Fig. 5B). It is possible that in the absence of EsxN, other ESX-5 substrates like PPE41 are more efficiently secreted. Both EspG_5_ and EccB_5_ were also produced at similar levels in the Δ*pstA1*Δ*esxN* double mutant compared to the Δ*pstA1* mutant (Fig. 5B). The Δ*pstA1*Δ*esxN* mutant hyper-secreted PPE41, like the Δ*pstA1* mutant (Fig. 5B). The ModD and GroEL2 controls demonstrate equivalent loading of culture filtrate and cell lysate fractions, respectively (Fig. 5B). Overall, our data suggest that EsxN is not required for production or stability of the ESX-5 components EccB_5_ or EspG_5_ and that EsxN and PPE41 are secreted independently.

To investigate if deletion of *esxN* could reverse attenuation of Δ*pstA1* mutant, like the Δ*BS* mutation, we infected Irgm1^-/-^, NOS2^-/-^ and C57BL/6 mice via the aerosol route with ∼100 CFU the Δ*pstA1*Δ*esxN* mutant. In contrast to the Δ*pstA1*Δ*BS* mutant that replicated progressively in the lungs of Irgm1^-/-^ mice (Fig. 4A), replication of the Δ*pstA1*Δ*esxN* mutant was well controlled in Irgm1^-/-^ mice (Fig. 5C). There was no significant difference in CFU recovered from the lungs of Δ*pstA1*- and Δ*pstA1*Δ*esxN*-infected Irgm1^-/-^ mice at any time point (Fig. 5C). The Δ*pstA1*Δ*esxN* mutant remained attenuated in NOS2^-/-^ mice and during the chronic phase of infection in C57BL/6 mice (Fig. 5D and 5E), similar to the Δ*pstA1*Δ*BS* mutant. Bacterial burdens in the lungs were not significantly different in either NOS2^-/-^ mice or C57BL/6 mice infected with the Δ*pstA1* mutant or the Δ*pstA1*Δ*esxN* mutant at any time point (Figs. 5D and 5E). These data suggest that ESX-5 secreted factors other than EsxN contribute to the attenuation of the Δ*pstA1* mutant in Irgm1^-/-^ mice.

## DISCUSSION

We previously demonstrated that the virulence-associated ESX-5 secretion system is regulated at the transcriptional level by the Pst/SenX3-RegX3 system that stimulates ESX-5 secretion in response to P_i_ limitation. By precisely defining the RegX3 binding site in the *esx-5* locus and creating targeted mutations that specifically disrupt RegX3-mediated regulation of ESX-5 secretion, we show here that regulation of ESX-5 secretion contributes to *M. tuberculosis* pathogenesis. Our data suggest that the Δ*pstA1* mutant is attenuated in Irgm1^-/-^ mice due to hyper-secretion of ESX-5 substrates caused by constitutive activation of RegX3. Our data further suggest that this attenuation is caused by ESX-5 substrates other than EsxN. Our results are consistent with a recent report demonstrating that secretion of the PE_PGRS subfamily of PE proteins, which are likely ESX-5 substrates, is associated with reduced *M. tuberculosis* virulence in Balb/c mice (20, 41). We conclude that *M. tuberculosis* requires precise regulation of ESX-5 secretion during infection for pathogenesis and that ESX-5 substrates other than EsxN play a direct role in the interaction with the host.

While our previous work demonstrated that RegX3 directly controls ESX-5 secretion at the transcriptional level and defined a region 5’ of the *pe19* gene to which RegX3 binds (14), the precise binding site remained unknown. Here we identified a RegX3 binding site sequence at −128 to −102 bp relative to the *pe19* start codon that consists of three imperfect direct repeats. We further demonstrated that the two 3’ direct repeats and 5 bp spacer with the sequence 5’-GGTGCcaactGGTGA-3’ are necessary for RegX3 binding *in vitro* and transcriptional regulation of *esx-5* genes *in vivo.* In this respect, RegX3 acts similarly to the *Escherichia coli* PhoB response regulator that also responds to P_i_ limitation by binding to direct repeat sequences (*pho* boxes) in the promoters of regulated genes (42). The RegX3 binding site sequence in the *esx-*5 locus is upstream of a transcriptional start site that was mapped at −38 relative to the *pe19* start codon (43), consistent with RegX3 acting as a transcriptional activator of *esx-5* genes. By creating mutations in or deleting this RegX3 binding site sequence on the *M. tuberculosis* chromosome, we demonstrate that regulation of ESX-5 secretion by RegX3 in response to P_i_ availability requires this sequence.

Attenuation of the Δ*pstA1* mutant specifically in Irgm1^-/-^ mice was almost completely suppressed by deletion of the RegX3 binding site in the *esx-5* locus, suggesting that the Δ*pstA1* mutant is attenuated in these mice due to constitutive ESX-5 secretion. These data also suggest that the Δ*pstA1* mutant is sensitive to some host factor other than Irgm1 due to constitutive ESX-5 secretion. Irgm1 and NOS2 act independently to control *M. tuberculosis* replication (29), so it is possible that constitutive ESX-5 secretion causes increased susceptibility of the Δ*pstA1* mutant to NOS2-generated nitrosative stress. Alternatively, the Δ*pstA1* mutant may fail to induce the generalized leukopenia that is typically observed in infected Irgm1^-/-^ mice (32), leading to improved control of the infection. In Irgm1^-/-^ mice, IFN-γ produced in response to infection causes the lymphatic collapse by stimulating autophagic death of effector T cells (31). Effector T cells may more efficiently recognize and control replication of the Δ*pstA1* mutant due to its constitutive secretion of antigenic ESX-5 substrates, so that T cell containment of infection occurs despite reduced T cell abundance. The Δ*pstA1* mutant could also interfere with IFN-γ production or signaling due to constitutive ESX-5 secretion. Manipulation of cytokine responses is a plausible explanation considering that ESX-5 has previously been implicated in activating the inflammasome and triggering IL-1β production by infected cells (18, 19). Finally, the susceptibility of Irgm1^-/-^ mice to infection with intracellular pathogens can be reversed by deletion of a second IFN-γ regulated GTPase Irgm3 (44). In Irgm1^-/-^ cells, mislocalization of effector immunity-related GTPases (IRGs) causes damage to lysosomes, but in cells lacking Irgm1 and Irgm3 the effector IRGs localize to lipid droplets and damage to lysosomes is prevented (45). It is possible that an ESX-5 secreted protein or proteins interferes with the function of either Irgm3 or the effector GTPases to prevent lysosomal damage and enable Irgm1^-/-^ mice to contain replication of the Δ*pstA1* mutant. We intend to explore these ideas in our future studies.

While constitutive ESX-5 secretion attenuates the Δ*pstA1* mutant in Irgm1^-/-^ mice, the ESX-5 substrates responsible for this phenotype remain to be determined. Our data suggest that attenuation is not caused by hyper-secretion of EsxN since the Δ*pstA1*Δ*esxN* mutant remained attenuated in Irgm1^-/-^ mice. It is possible that one or more of the EsxN paralogs (EsxI, EsxL, EsxO, or EsxV) plays some role in this process. We could still detect secretion of one or more of these proteins from the Δ*pstA1*Δ*esxN* mutant using our EsxN anti-serum. However, secretion of all EsxN paralogs was undetectable in the Δ*pstA1*Δ*BS* mutant, suggesting that decreased production of ESX-5 core components reduces secretion of all EsxN paralogs. Our future plans include deleting genes encoding each of the EsxN paralogs individually and in combination to determine whether these proteins collectively influence pathogenesis. Alternatively, PE and/or PPE proteins secreted via ESX-5 may play a role in attenuation of the Δ*pstA1* mutant. PE and PPE proteins are strongly immunogenic in mice in a manner dependent on secretion via ESX-5 (21). In addition, some PE and PPE proteins can directly manipulate the functions of host cells (46-48) and, as discussed above, secretion of the PE_PGRS subfamily in particular has previously been associated with reduced virulence (41). We are currently working to define the *M. tuberculosis* ESX-5 secretome using strains we developed that conditionally express the ESX-5 core component EccD_5_ (33) and will explore the potential of these secreted substrates to influence pathogenesis.

Although our data indicate that ESX-5 hyper-secretion causes attenuation of the Δ*pstA1* in Irgm1^-/-^ mice, aberrant ESX-5 secretion does not contribute substantially to the chronic phase persistence defect of the Δ*pstA1* mutant in C57BL/6 mice. We recently described that the Δ*pstA1* mutant also exhibits increased release of membrane vesicles (MV) derived from the inner membrane that contain immune-modulatory lipoproteins and lipoglycans (33, 49). Importantly, increased MV release by the Δ*pstA1* mutant was independent of ESX-5 secretion system activity (33). We speculate that aberrant MV production could also contribute to attenuation of the Δ*pstA1* mutant. The MV-associated lipoprotein LpqH (also known as the 19kDa lipoprotein) is a potent TLR2 ligand and signaling through this pathway causes pleiotropic effects on the innate immune system that include promoting the production of the pro-inflammatory cytokines IL-1β, IL-12p40, and TNFα (50), reducing surface MHC class II expression on macrophages (50-52), and inducing host cell apoptosis and nitric oxide-independent antimicrobial activity (49, 53, 54). Additionally, LpqH contributes to CD4^+^ T cell activation (55). It is unclear from these studies if increased release of LpqH would be beneficial or detrimental to bacterial survival and pathogenesis. We are actively exploring the mechanism of enhanced MV release by the Δ*pstA1* mutant to determine the importance of regulated MV production in *M. tuberculosis* pathogenesis.

While constitutive activation of ESX-5 secretion contributes to attenuation of the Δ*pstA1* mutant, regulation of ESX-5 secretion by RegX3 appears to play only a minor role in *M. tuberculosis* pathogenesis. *regX3* mutants are attenuated during chronic infection of C57BL/6 mice (24, 56), but the Δ*BS* mutant that we constructed, which fails to induce transcription of *esx-5* genes or secretion of ESX-5 substrates in response to P_i_ limitation *in vitro*, was only modestly attenuated during the acute phase of infection and persisted normally in the chronic phase of infection. Our data suggest that other regulatory targets of RegX3 besides ESX-5 influence *M. tuberculosis* persistence and that other regulators may contribute more substantially to controlling ESX-5 activity during infection. Indeed several transcription factors have been reported to bind within the *esx-5* locus and induce transcription of *esx-5* genes (34, 57). It is possible that one or more of these regulators plays an important role in controlling ESX-5 secretion during infection, which we plan to investigate in our future studies.

## MATERIALS AND METHODS

### Bacterial strains and culture conditions

*M. tuberculosis* Erdman and the derivative Δ*pstA1*, Δ*regX3*, and Δ*pstA1*Δ*regX3* mutant strains were previously described (24). Construction of strains harboring mutations in the *esx-5* RegX3 binding site sequence is described below. Bacterial cultures were grown at 37°C with aeration in Middlebrook 7H9 liquid medium (Difco) supplemented with albumin-dextrose-saline (ADS), 0.5% glycerol and 0.1% Tween-80 or on Middlebrook 7H10 agar medium (Difco) supplemented with 10% Middlebrook oleic acid-albumin-dextrose-catalase (OADC, BD Biosciences) and 0.5% glycerol, unless otherwise noted. Sauton’s medium (3.67 mM KH_2_PO_4_, 2 mM MgSO_4_-7H_2_O, 9.5 mM citric acid, 0.19 mM ammonium iron (III) citrate, 26.64 mM L-asparagine, 6% glycerol, 0.01% ZnSO_4_, pH 7.4) or P_i_-limited Sauton’s medium (Sauton’s containing 2.5 μM KH_2_PO_4_, buffered with 50 mM MOPS, pH 7.4) were used to grow cultures for protein isolation. P_i_-free 7H9 medium was prepared as previously described (24). Frozen stocks were prepared by growing liquid cultures to mid-exponential phase (OD_600_ 0.8-1.0) in complete 7H9 medium, then adding glycerol to 15% final concentration, and storing 1 ml aliquots at −80°C.

### Cloning

Constructs for deleting *esxN* or introducing mutations in the *esx-5* locus RegX3 binding site were generated in the pJG1100 allelic exchange vector, which contains the *aph* (kanamycin resistance), *hyg* (hygromycin resistance), and *sacB* (sucrose sensitivity) markers (58). Genomic regions ∼800 bp 5’ and 3’ of the sequence to be mutated were PCR amplified from the *M. tuberculosis* Erdman genome using the primers in Table S2. Forward primers to amplify the 5’ region were designed with a PacI restriction site; reverse primers to amplify the 3’ region were designed with an AscI restriction site. For deletion of *esxN,* the reverse primer to amplify the 5’ regions and the forward primer to amplify the 3’ region were designed with AvrII restriction sites in-frame with the start and stop codons, respectively. Resulting PCR products were cloned in pCR2.1 (Invitrogen) and sequenced. The 5’ and 3’ regions were removed from pCR2.1 by restriction with PacI/AvrII and AvrII/AscI, respectively, and ligated with pJG1100 digested with PacI/AscI to generate the in-frame Δ*esxN* deletion construct. For the *esx-5* RegX3 binding site mutations, the forward and reverse primers for amplifying the 3’ and 5’ regions of homology, respectively, contained the mutation to be introduced and were designed with overlapping sequence at the 5’ ends to allow PCR products to be joined by overlap extension PCR (59) before cloning in pCR2.1. Sequence-confirmed binding site mutation constructs were removed from pCR2.1 by restriction with PacI/AscI and ligated to similarly digested pJG1100.

### Strain construction

*M. tuberculosis* strains harboring the Δ*esxN* deletion or *esx-5* RegX3 binding site mutations were generated by two-step allelic exchange, as previously described (24). Integration of the pJG1100 construct at the correct location was confirmed by colony PCR on heat-inactivated cell lysates using the primer pairs for detection of the 5’ and 3’ homologous recombination (Tables S2 & S3). Clones with the plasmid integrated were grown without antibiotics, diluted and plated on 7H10 containing 2% sucrose for counter-selection of the pJG1100 vector. Sucrose resistant isolates were screened by colony PCR on heat-inactivated cell lysates using primers for the detection of the deletion or mutation (Tables S2 & S3). The *esx-5* RegX3 binding site mutations were verified by Sanger sequencing of the resulting PCR products.

### Purification of His_6_-RegX3

Recombinant His_6_-RegX3 was expressed and purified from *Eschericia coli* BL21 (DE3) containing pET28b+::*regX3* by affinity chromatography using Ni-NTA agarose (Qiagen) as previously described (14).

### Electrophoretic mobility shift assays

Double-stranded DNA probes were PCR amplified using *M. tuberculosis* Erdman genomic DNA as template and appropriate primers (Table S4). Probes were labeled with the DIG Gel Shift Kit, 2^nd^ Generation (Roche), following the manufacturer’s protocols. Binding reactions with 0.5 ng of DIG-labeled probe, binding buffer (Roche), poly[d(I-C)], poly L-lysine, and 0.5 μg purified His_6_-RegX3 in 20 μl total volume were incubated at room temperature for 15 min. Binding reactions including a 400-fold excess of unlabeled competitor (200 ng) were incubated for 15 min at room temperature prior to adding the DIG-labeled probe, then incubated an additional 15 min. DNA-protein complexes were resolved by electrophoresis on 5% native polyacrylamide gels, transferred and UV-crosslinked to nylon membranes (Roche). Membranes were washed with wash buffer (DIG wash and block buffer set, Roche), blocked for 30 min in blocking solution (Roche) and incubated with anti-DIG-AP antibodies (Roche) at a 1:10,000 dilution for 30 min at room temperature. Labeled probes were detected using CDP-Star ready-to-use substrate (Roche). Membranes were exposed to film (Blue lite autrorad film, Genemate) and developed using a film processor (Konica, SRX-101A).

### Quantitative RT-PCR

To measure gene expression in P_i_-rich conditions, bacteria were grown in complete Middlebrook 7H9 medium to mid-exponential phase (OD_600_ 0.4-0.6). To test induction of gene expression during P_i_ starvation, cultures were grown in 7H9 to mid-exponential phase (OD_600_ 0.4-0.6), washed twice and resuspended at OD_600_ 0.2 in P_i_-free 7H9, and then grown at 37°C with aeration for 24 hr. Bacteria were collected by centrifugation (3700 x *g*, 10 min, 4°C). Total RNA was extracted using TRIzol (Invitrogen, CA) with 0.1% polyacryl carrier (Molecular Research Center, Inc) by bead beating with 0.1 mm zirconia beads (BioSpec Products). Equivalent amounts of total RNA were treated with Turbo DNase (Invitrogen) and converted to cDNA using the Transcriptor First Strand cDNA Synthesis Kit (Roche) with random hexamer primers and the following parameters: 10 min at 25°C (annealing of primers), 60 min at 50°C (elongation), and 5 min at 85°C (heat inactivation of reverse transcriptase). cDNA was stored at −20°C.

Quantitative PCR primers to amplify internal regions of the genes of interest (*pe19, esxN, espG*_*5*_, *eccD*_*5*_, *udgA, mgtA,* and *sigA*) were designed with similar annealing temperatures (58-60°C) using either Primer Express software (Applied Biosystems) or ProbeFinder Assay Design software (Roche) and are listed in Table S5. Quantitative RT-PCR reactions were prepared using 2x SYBR Green master mix (Roche), 2.5 µM each primer and 1 µl cDNA and run on a LightCycler 480 (Roche) using the following cycle parameters: 95°C for 10 min; 45 cycles of 95°C for 10s, 60°C for 20s, and 72°C for 20s with data collected once per cycle during the extension phase; and one cycle of 95°C for 5s, 65°C for 1m, 97°C with a ramp rate of 0.11°C/s for generation of melting curves. Cycle threshold values (C_p_, Roche nomenclature) were converted to copy numbers using standard curves for each gene generated using genomic DNA. Gene copy numbers were normalized to *sigA*.

### Antisera production

Rabbit polyclonal antisera against EccD_5_, EsxN and PPE41 were previously described (33, 60). Synthetic antigenic peptides (EccB_5_ 489-506, EHDTLPMDMTPAELVVPK; EspG_5_ 283-300, KTVLDTLPYGEWKTHSRV) that were identified with Antigen Profiler and conjugated to keyhole limpet hemocyanin (KLH) were used with TiterMax Gold adjuvant (Sigma) to raise polyclonal antisera against EccB_5_ and EspG_5_ in rabbits (Pierce Custom Antibodies, Thermo Scientific).

### Protein preparation for immunoblots

*M. tuberculosis* cultures were grown at 37ºC with aeration in Sauton’s medium or P_i_-limited (2.5 μM P_i_) Sauton’s medium for five days as previously described (14) prior to protein isolation. Bacteria were collected by centrifugation (4700 x *g*, 15 min, 4°C). Culture supernatants were filter sterilized as previously described (14) and Complete EDTA-free protease inhibitor tablets (Roche) were added. Supernatants were concentrated roughly 25-fold by centrifugation (2400 x *g*, 4°C) using VivaSpin 5 kDa molecular weight cut-off spin columns (Sartorius). Whole cell lysates were prepared by bead beating with 0.1 mm zirconia beads (BioSpec Products) in PBS containing Complete EDTA-free protease inhibitors (Roche) and lysates were clarified by centrifugation as previously described (14). Cell lysates were passaged through a Nanosep MF column with a 0.22 µm filter (Pall Life Sciences) by centrifugation (14000 × *g*, 3 min, 4°C) to remove any remaining intact cells. Total protein concentration in each sample was quantified using the Pierce BCA Protein Concentration Assay kit (Thermo Scientific). Proteins were stored at 4°C for immediate use, or at −80°C with glycerol at 15% final concentration.

### Western blotting

Culture filtrate or whole cell lysate proteins were separated by sodium dodecyl sulfate polyacrylamide gel electrophoresis (SDS-PAGE) on Mini-PROTEAN TGX Any kD gels (Bio-Rad) and transferred to nitrocellulose membranes (Whatman) by electrophoresis. Proteins were detected by Western blotting as previously described (14) using primary anti-sera at the following diutions: rabbit α-EsxN 1:1000; rabbit α-EspG_5_ 1:1000; rabbit α-EccB_5_ 1:1000; rabbit α-EccD_5_ 1:1000; rabbit α-PPE41 1:1000; rabbit α-ModD 1:25000; mouse α-GroEL2 1:1,000. Appropriate secondary antibodies (either goat-anti-rabbit or rabbit-anti-mouse conjugated to HRP, Sigma) and SuperSignal West Pico substrate (Thermo Scientific) were used to detect reactive bands. Blots were imaged on an Odyssey Fc Imaging System (LI-Cor) and protein abundance was analyzed using ImageStudio software (LI-Cor).

### Mouse infections

Female C57BL/6J and NOS2^-/-^ mice 6-8 weeks of age were purchased from Jackson Laboratories. Irgm1^-/-^ mice were bred under specific-pathogen-free conditions at the University of Minnesota Research Animal Resources. Mice were infected with ∼100 CFU using an Inhalation Exposure System (GlasCol) as previously described (36). Infected mice were euthanized with CO_2_ overdose. Bacterial CFU were enumerated by plating serial dilutions of lung homogenates on complete Middlebrook 7H10 agar containing 100 μg/ml cyclohexamide and counting CFU after 3-4 weeks of incubation at 37°C. All animal protocols were reviewed and approved by University of Minnesota Institutional Animal Care and Use committee and were conducted in accordance with recommendations in the National Institutes of Health *Guide for the Care and Use of Laboratory Animals* (61).

## ACKNOWLEDGEMENTS

We thank Alyssa Brokaw and Leanne Zhang for expert technical assistance with animal experiments, and the staff of the University of Minnesota BSL-3/ABSL-3 core facility. Antisera against GroEL2 (monoclonal clone IT-70, cat. no. NR-13657) and ModD (polyclonal anti-Mpt32, cat. no. NR-13807) were obtained from BEI Resources, NIAID, NIH. This work was supported by an NIH Director’s New Innovator Award, DP2AI112245 (A.D.T.), start-up funding from the University of Minnesota (A.D.T.), and the Dennis W. Watson Fellowship (D.W.W.).

